# Generation of parasympathetic neurons from hiPSC that reproduce the electrophysiological properties of native neurons and modulate the activity of hiPSC-atrial cardiomyocytes

**DOI:** 10.1101/2025.10.23.684157

**Authors:** Alison M. Thomas, Isabella Noelle Chiong, Joana F Neves, Andrew Tinker, Franziska Denk, Laura Fedele

**Affiliations:** Wolfson Sensory Pain and Regeneration Centre, King’s College London, London, SE1 1UL, United Kingdom; Clinical Pharmacology & Precision Medicine, William Harvey Research Institute, Barts and the London School of Medicine and Dentistry, Queen Mary University of London; Centre for Host Microbiome Interactions, King’s College London

## Abstract

Peripheral parasympathetic ganglia lie adjacent to their target organs; their dysfunction contributes to diseases like atrial fibrillation. Due to their challenging accessibility for primary culture, we provide a step-by-step protocol to generate parasympathetic neurons from human induced pluripotent stem cells (hiPSC). They reproduce numerous native physiological features: cholinergic and autonomic markers, functional nicotinic receptors and electrophysiological properties. We also describe a co-culture system with hiPSC-atrial cardiomyocytes allowing neuromodulation of cardiomyocyte activity and offering a platform for drug discovery and disease modelling.

**Highlights:**

- Steps to generate parasympathetic neurons from human induced pluripotent stem cells
- Guidance of the functional assessment of the neurons (whole-cell patch clamp and Ca^2+^ imaging)
- Instructions on how to co-culture these neurons with hiPSC-derived atrial cardiomyocytes
- Guidance of the functional assessment of the neuronal modulation of cardiomyocyte activity

**Graphical abstract:** 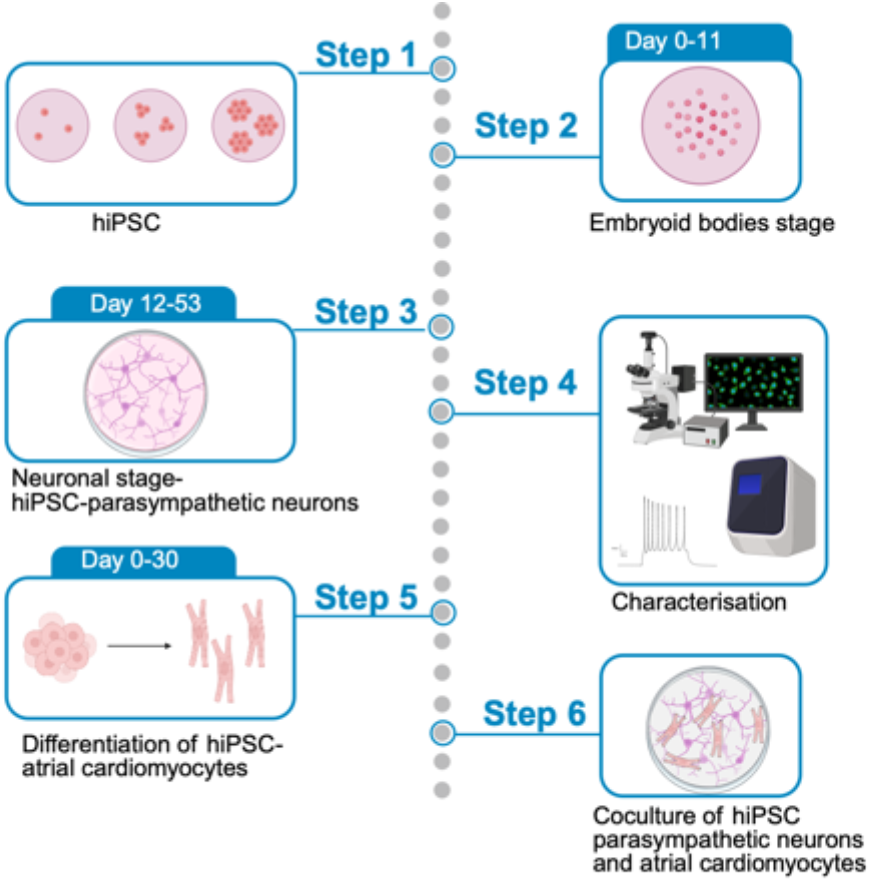

## Innovation

This protocol was adapted from a pioneering cranial motor neuron differentiation protocol provided by^1^. We added a maturation step and adapted their protocol for manual handling, substituting the V-shaped plates and the Bravo Automated Liquid Handling platform with affordable non-tissue culture treated dishes. In contrast to other protocols^2^, our protocol does not require batch-testing of growth factors, like BMP4, making it easier for new users. For the first time, we also characterized the cells for their electrophysiological and functional properties and beyond markers that distinguish them from spinal motor neurons. We demonstrated physiologically that our parasympathetic neurons resemble their native counterparts in several important and novel ways: they respond to nicotine, from day 39, have a resting membrane potential of –65mV, and can be grouped into those that generate phasic versus tonic action potentials. We cannot say for sure if they are preganglionic or postganglionic parasympathetic neurons, however their expression of the postganglionic parasympathetic marker HMX3^3,4^, not expressed in undifferentiated cells, would suggest some postganglionic lineage. Finally, we demonstrated that they can form a functional connection with hiPSC-atrial cardiomyocytes: nicotine application to our coculture (hiPSC-parasympathetic neurons with hiPSC-atrial cardiomyocytes) reduced spontaneous Ca^2+^ transients in cardiomyocytes.

## Before you begin

### Thawing and maintenance of human induced Pluripotent stem cells (hiPSC) before differentiating them

#### Timing: 2 weeks

We tested the protocol using two hiPSC lines (one male and one female) that required two distinct methods of matrix coating. The Kute4 line grew better in vitronectin coated plates, whereas the UKB line in Geltrex coated plates. The differentiation protocol worked for both cell lines; differences were only in the matrix used to grow hiPSC and how they were passaged for maintenance.

1. Coating plates with Vitronectin *Dilute vitronectin in room temperature Dulbecco phosphate base solution (DPBS) without Magnesium or Calcium, for a final solution of 10μg/mL (80μL in 2mL)*
  a. For a 6-well plate, coat 1mL per well. One well is enough per hiPSC vial
  b. Leave the well coated for 1-2hours under the tissue culture hood at room temperature
  c. Remove the solution just before seeding cells **NOTE**: To avoid freezing/thawing cycles, store Vitronectin in the freezer in 40*μ*L or 80*μ*L aliquots
2. Coating plates with Geltrex Dilute Geltrex (1:50) in cold basal Stemflex without any supplement
  a. For a 6-well plate, coat 1mL per well. One well is enough per hiPSC vial
  b. Leave the well coated for 1 hour under the tissue culture hood at room temperature
  c. Remove the solution just before seeding cells **CRITICAL:** Geltrex is temperature dependent; to avoid premature gelling, thaw the frozen vial in cold media and store in 50-100*μ*L aliquots. The timing of matrix coating is critical, especially for Geltrex: ensure to coat the well for no longer than 1h 30min, longer coating can affect gelling and compromise cell adherence.
3. Thawing cells
  a. Prepare a 15ml falcon tube with 5mL of Stemflex media (Table 1)
  b. Thaw hiPSC vial in a water bath (37°C) swirling the vial, until nearly fully thawed (i.e. until only a small ice ball is visible)
  c. Transfer the cells into the tube with media
  d. Centrifuge at 280g for 5min
  e. Remove and discard the supernatant without disrupting the pellet
  f. Gently resuspend the pellet with 1mL Stemflex complete media (Table 1), supplemented with 1*μ*M Rock inhibitor
  g. Remove coating matrix
  h. Plate cells
  i. Wash the tube with an additional 500*μ*L of Stemflex media (Table 1) with 1*μ*M Rock inhibitor and add to well
  j. Distribute the cells evenly, swirling the plate with a circular motion
4. Media change
  a. Change full media (1.5mL per well of a 6-well plate) 24h after plating; then every 48h for maintenance or, e.g. to avoid weekend work, double feed (i.e. 3mL per well for a 6-well plate)
5. Passaging cells (vitronectin coated wells)
  a. Passage cells when they are 70% confluent
  b. Coat wells 45-50 min before starting
  c. Wash cells twice with PBS (without Mg^2+^/Ca^2+^)
  d. Incubate with versene for 4 minutes (37°C, 5% CO_2_)
  e. Remove versene, add 1mL Stemflex media and detach the cells
  f. Remove vitronectin
  g. Plate cells at the desired dilution, topping up to a total of 1.5mL media per well
  h. Distribute the cells evenly swirling the plate with a circular motion
6. Passaging cells (Geltrex coated wells)
  a. Passage cells when they are 50-60% confluent
  b. Coat wells 45-50 min before starting
  c. Wash cells twice with PBS (without Mg^2+^/Ca^2+^)
  d. Incubate with 0.5mM EDTA (in PBS without Mg^2+^/Ca^2+^) at room temperature (under the hood) for 5 minutes
  e. Remove EDTA, add 1mL Stemflex media and detach the cells
  f. Remove Geltrex
  g. Plate cells at the desired dilution, topping up to a total of 1.5mL media per well
  h. Distribute the cells evenly swirling the plate with a circular motion

**Table 1:** Stemflex media.

|  | Final concentration | Amount |
| --- | --- | --- |
| Stemflex basal media | 89% | 44.5mL |
| Stemflex supplement | 10% | 5mL |
| Antimycotic-antibiotic | 1% | 0.5mL |

Note on storage: store wrapped in foil (light sensitive!) at 4°C for up to two weeks.

**CRITICAL**: Passage cells at least two times before starting a differentiation protocol. Make sure to start with healthy hiPSC colonies (see Troubleshooting, Problem 1)

**NOTE**: For maintenance, we usually split the hiPSC at 1:10/1:20. When passaging to start a differentiation, we ensure that the colonies do not merge, removing merged colonies with a p10 tip (illustrated in Figure 6). We found that passaging cells at a 1:20 dilution was the optimal split to have non-overlapping colonies for both cell lines; 4-5 days of expansion was the optimal time before starting a differentiation. However, these dilution factors and timings might vary, depending on the iPSC line and the matrix it grows on.

### Embryoid bodies stage

#### Timing: 10 days

Per batch we use two wells of a 6-well plate, which yields a total of 5-7.5 million cells (70-100 coverslips) on day 12. For the embryoid stage, we combine the wells in a non-tissue culture coated 4cm dish. An overview of the protocol is shown in Figure 1.

**Figure 1.**
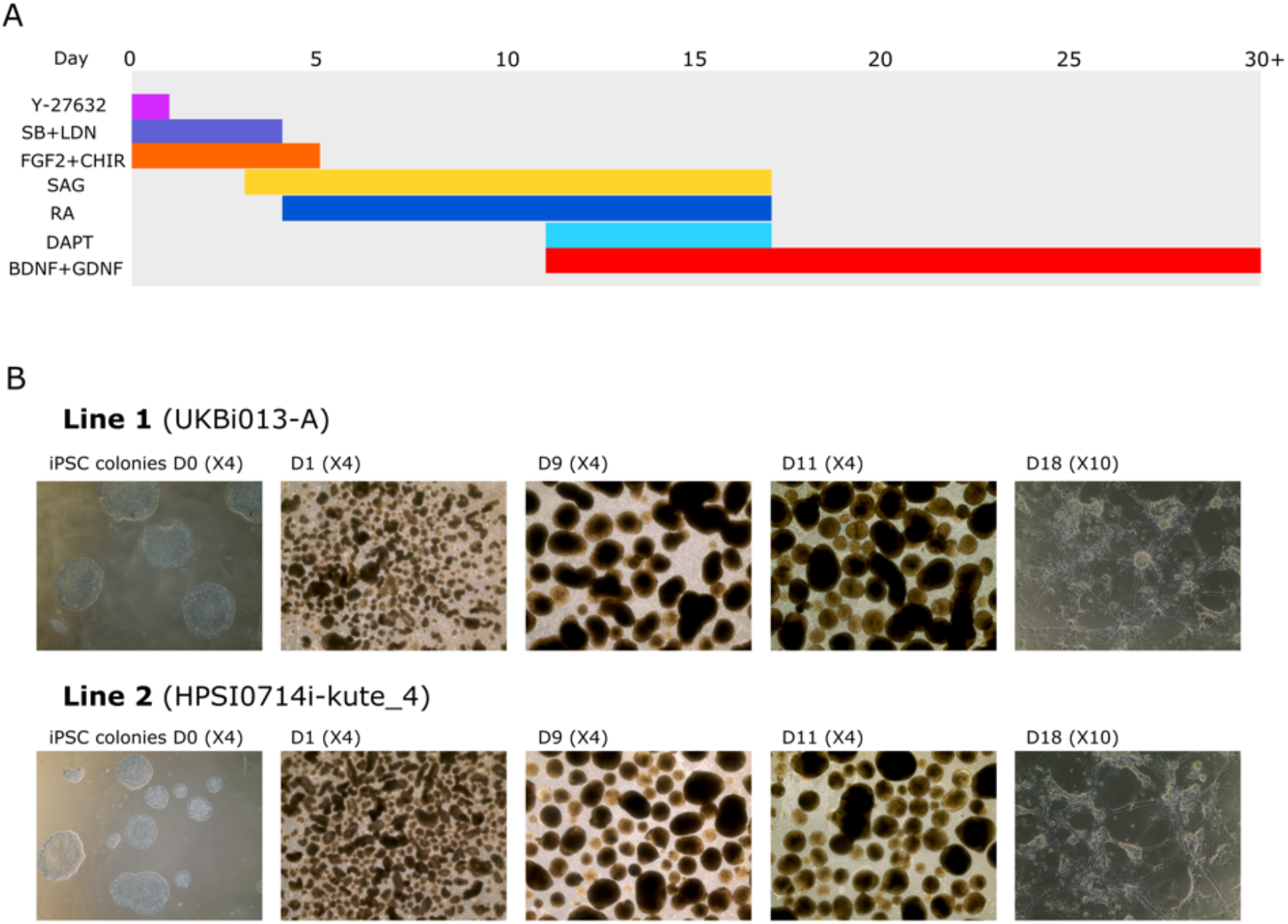
Differentiation of parasympathetic neurons from hiPSC. (A) Overview of the protocol. (B) and (C) Phase contrast images from day 0 to day 18 using the two hiPSC lines.

7. Day 0
  a. Wash colonies with HBSS (without Ca^2+^/Mg^2+^) or PBS (without Ca^2+^/Mg^2+^)
  b. Incubate with 1:100 collagenase IV solution (50000U/mL stock) (Table 2) at 37°C for 1h. Use 1mL of collagenase solution per well
  c. Inactivate the solution by adding 2mL of DMEM F12/Neurobasal (1mL DMEM F12, 1 mL Neurobasal) per well.
  d. Transfer cells in solution to a 15mL tube, wash wells with an additional 2mL and add to the tube
  e. Allow the hiPSC colonies to form a pellet in the 15mL tube by waiting for a couple of minutes or centrifuging at 300g for 2min
  f. Remove supernatant
  g. Transfer 4mL of **Day 0 complete media (Table 4)** to a 6cm non-TC treated dish
  h. Resuspend the pellet gently with 500*μ*L of the above media from the 6cm dish
  i. Pipette up and down 2-3 times (P1000) and transfer colonies to the dish

**Table 2:** Collagenase solution.

|  | Final Concentration | Amount |
| --- | --- | --- |
| Collagenase (Type IV) | 1:100 | 10μL |
| DMEM F12 |  | 0.5mL |
| Neurobasal |  | 0.5mL |

Note on storage: prepare collagenase aliquots (at 50000U/ml in HBSS with Ca<sup>2+</sup> and Mg<sup>2+</sup>). Store in 10uL at -20°C, dilute in warm media just before usage. NB collagenase is light-sensitive (wrap in foil when thawing).

**Table 3:** Parasympathetic neuron media.

|  | Concentration | Amount |
| --- | --- | --- |
| DMEM F12 | 47.95 % | 23.975mL |
| Neurobasal | 47.95 % | 23.975mL |
| Pen-strep (or Anti/anti) | 1% | 0.5mL |
| b-mercaptoethanol (50mM) | 50μM | 0.05mL |
| N2 | 1% | 0.5mL |
| B27 | 2% | 1mL |

**NOTE:** Collagenase incubation lifts the hiPSC colonies from the wells, this allows easier formation of embryoid bodies as the cells are already organized in a circular shape.

**CRITICAL:** Do not triturate colonies too much; you can also use a P1000 cut tip. Do not transfer colonies directly on the dish without media, as it can affect cell health.

8. Day 1-11
  a. Tilt the dish, remove as much media as possible without disrupting the embryoid bodies
  b. Supplement with appropriate media as outlined in Table 4

**Table 4:** Media composition for the embryoid bodies/neurosphere stage – the small molecules and inhibitors are diluted in parasympathetic media (Table 3)

| Day | Concentration | Dilution in 4mL |
| --- | --- | --- |
| 0 | 5μM Y-27632<br>40μM SB431542 | 2μL Y-27632 (Ri)<br>8μL SB431542 |
|  | 0,2µM LDN193189<br>10ng/ml FGF2<br>1µM Chir-99021<br>0.5µM Ascorbic Acid | 4µL LDN193189<br>4µL FGF2<br>0.65µL Chir-99021<br>0.4µL Ascorbic Acid |
| 1 | 40µM SB431542<br>0,2µM LDN193189<br>10ng/ml FGF2<br>1µM Chir-99021<br>0.5 µM Ascorbic Acid | 8µL SB431542<br>4µL LDN193189<br>4µL FGF2<br>0.65µL Chir-99021<br>0.4µL Ascorbic Acid |
| 2 | 40µM SB431542<br>0.2µM LDN193189<br>10ng/ml FGF2<br>1µM Chir-99021<br>0.5µM Ascorbic Acid | 8µL SB431542<br>4µL LDN193189<br>4µL FGF2<br>0.65µL Chir-99021<br>0.4µL Ascorbic Acid |
| 3 | 40 µM SB431542 (Tocris)<br>0,2 µM LDN193189 (Miltényi)<br>10ng/ml FGF2<br>1µM Chir-99021<br>500nM SAG<br>0.5µM Ascorbic Acid (Sigma) | 8µL SB431542<br>4µL LDN193189<br>4µL FGF2<br>0.65µL Chir-99021<br>4µL SAG<br>0.4µL Ascorbic Acid |
| 4 | 10ng/ml FGF2<br>1µM Chir-99021<br>500nM SAG<br>10nM Retinoic Acid<br>0.5µM Ascorbic Acid (Sigma) | 4µL FGF2<br>0.65µL Chir-99021<br>4µL SAG<br>0.4µL Retinoic Acid<br>0.4µL Ascorbic Acid |
| 5 | 500nM SAG<br>10nM Retinoic Acid<br>0.5µM Ascorbic Acid (Sigma) | 4µL SAG<br>0.4µL Retinoic Acid<br>0.4µL Ascorbic Acid |
| 7 | 500nM SAG<br>10nM Retinoic Acid<br>0.5µM Ascorbic Acid (Sigma) | 4µL SAG<br>0.4µL Retinoic Acid<br>0.4µL Ascorbic Acid |
| 9 | 500nM SAG<br>10nM Retinoic Acid<br>0.5µM Ascorbic Acid (Sigma) | 4µL SAG<br>0.4µL Retinoic Acid<br>0.4µL Ascorbic Acid |

NOTE: Dilute the small molecules inhibitors in media on the day. Change media on Day 0, 1,2,3,4,5,7 and 9. When changing media, tilt the dish, remove as much media as possible without disrupting the neurospheres.

**NOTE:** Change media only on Days 0, 1,2,3,4,5,7 and 9.

**CRITICAL:** Supplement the media with the correct small molecule inhibitor mixture for each appropriate day.

### Replating

#### Timing: 6 hours

9. Acid Treatment of coverslips
  a. Submerge 100 coverslips in acid solution (33ml 70% nitric acid +16.5ml of 34% HCl per bottle)
  b. under a fume hood
  c. Swirl every hour for 3-4 hours
  d. Wash them twice (or until no acid traces can be seen) with distilled water
  e. Store in 70% EtOH until use

**NOTE:** Acid treating coverslips reduces impurities and enhances Geltrex coating and cell adhesion. Coverslips can be acid treated up to one month before their usage. Once acid treated, we keep them in 80mL clean (autoclaved) glass bottles in 70%EtOH.

10. Preparation of coverslips in tissue culture plates
  a. Place coverslips in 24-well plates
  b. Wash them with PBS twice
  c. Remove PBS
  d. Using forceps, ensure that the coverslips are in the middle of the well
  e. Leave the lid of the plate open and allow coverslips to dry in the hood (around 20min)

**NOTE:** Ensuring that coverslips are in the middle of the well will make it easier to coat them with Geltrex and plate the cells. Ensure no contamination happens during the process.

11. Coating coverslips with Geltrex
  a. Dilute hESC-qualified Geltrex (1:50) in cold mixture of DMEM F12/Neurobasal media (1:1)
  b. Place a 60*μ*L bubble in the center of the coverslip

**CRITICAL**: Seeding cells on Geltrex coated dishes is a time-sensitive process, poor cell plating/seeding can result in cell death/lack of neurite formation (See Troubleshooting-Problem 2)

12. Dissociation of neurospheres
  a. Remove media from the 6cm dish without disrupting the neurospheres
  b. Collect the neurospheres with PBS (4mL) and transfer them into a 15mL tube
  c. Allow them to settle in the bottom, remove PBS and add 4mL PBS
  d. Remove PBS and incubate with 1mL TrypLE (37°C, 5%CO_2_), keeping the tube in the upright position with the pellet on the bottom of the tube
  e. Incubate at 37°C for 20min (time might vary according to hIPSC line). During the incubation, occasionally pipette up and down once (without triturating) to ensure the embryoid bodies are fully mixed in TrypLE. We normally pipette at minutes: 7, 10, 15, 18
  f. Following the 20min incubation: triturate the embryoid bodies by pipetting up and down 15-20 times, without introducing bubbles. The solution should become cloudy.
  g. Inactivate TrypLE with 3mL of basal media (50% DMEMF12/50% Neurobasal)
  h. Gently resuspend and pass through the cell suspension in a 70*μ*m cell strainer
  i. Count cells and resuspend at the required cell density in Day 11 complete media
  j. Just before plating, remove the Geltrex bubble
  k. Plate a 65*μ*L bubble per coverslip
  l. Wait 45-60min to allow cells to attach on the center of the coverslip
  m. Gently add 435*μ*L of complete Day 11 media

**NOTE:** Embryoid bodies are washed twice with PBS to avoid any media contamination that could inhibit TryplE. In our hands, 75,000 and 50,000 cells per coverslips both worked well.

**CRITICAL:** When incubating in TryplE, ensure that all the embryoid bodies are fully submerged.

### Neuron Stage

#### Timing: 20-30 days

13. Change media on Day 14, Day 17 (±1 day), as indicated in Table 5 and twice a week thereafter.

**Table 5:** Media composition for the neuronal stage – the small molecules and inhibitors are diluted in parasympathetic media.

|  | Concentration | Dilution in 40mL media |
| --- | --- | --- |
| 11 | 10µM Y-27632 (Ri)<br>500nM SAG<br>10nM Retinoic Acid<br>10µM DAPT | 40µL Y-27632 (Ri)<br>40µL SAG<br>4µL RA<br>20µL DAPT |
| | 20ng/mL BDNF<br>10ng/mL GDNF<br>0.5 μM Ascorbic Acid (Sigma) | 8μL BDNF<br>4μL GDNF<br>4μL Ascorbic acid |
| 14 | 500nM SAG<br>10nM Retinoic Acid<br>10μM DAPT<br>20ng/mL BDNF<br>10ng/mL GDNF<br>0.5 μM Ascorbic Acid (Sigma)<br>1/250 Geltrex (non-ESC) | 40μL SAG<br>4μL RA<br>20μL DAPT<br>8μL BDNF<br>4μL GDNF<br>4μL Ascorbic acid<br>160μL Geltrex (non-ESC) |
| 17 (±1 Day)><br>*change media twice a week;*<br>*add 1/250 Geltrex (nonesc)*<br>*once a week or every other*<br>*week* | 20ng/mL BDNF<br>10ng/mL GDNF<br>0.5μM Ascorbic Acid (Sigma) | 8μL BDNF<br>4μL GDNF<br>4μL Ascorbic acid |

**NOTE:** Perform full media changes, but very gently, do not let cells dry out. We never change more than 12 wells at a time. Geltrex is supplemented in the media (as detailed in Table 5) to avoid cells from detaching.

14. Quality control:
  a. At Day 14-17: Isolate RNA using the RNeasy Plus Microkit: combine 2-3 24-well coverslips, pipetting 350*μ*L of buffer RLT supplemented with beta-mercaptoethanol (1% v/v). Perform RT-qPCR for autonomic lineage markers and cholinergic markers (Figure 2). Collect RNA combining 2-3 of the 24-well coverslips closer to maturity stage (>Day 30) to further confirm their character, before using them for experiments.
  b. Fix coverslips at >Day 30 with 0.5mL 4% paraformaldehyde at room temperature for 10 min to test for PHOX2B and CHAT antibody staining.

**Figure 2.**
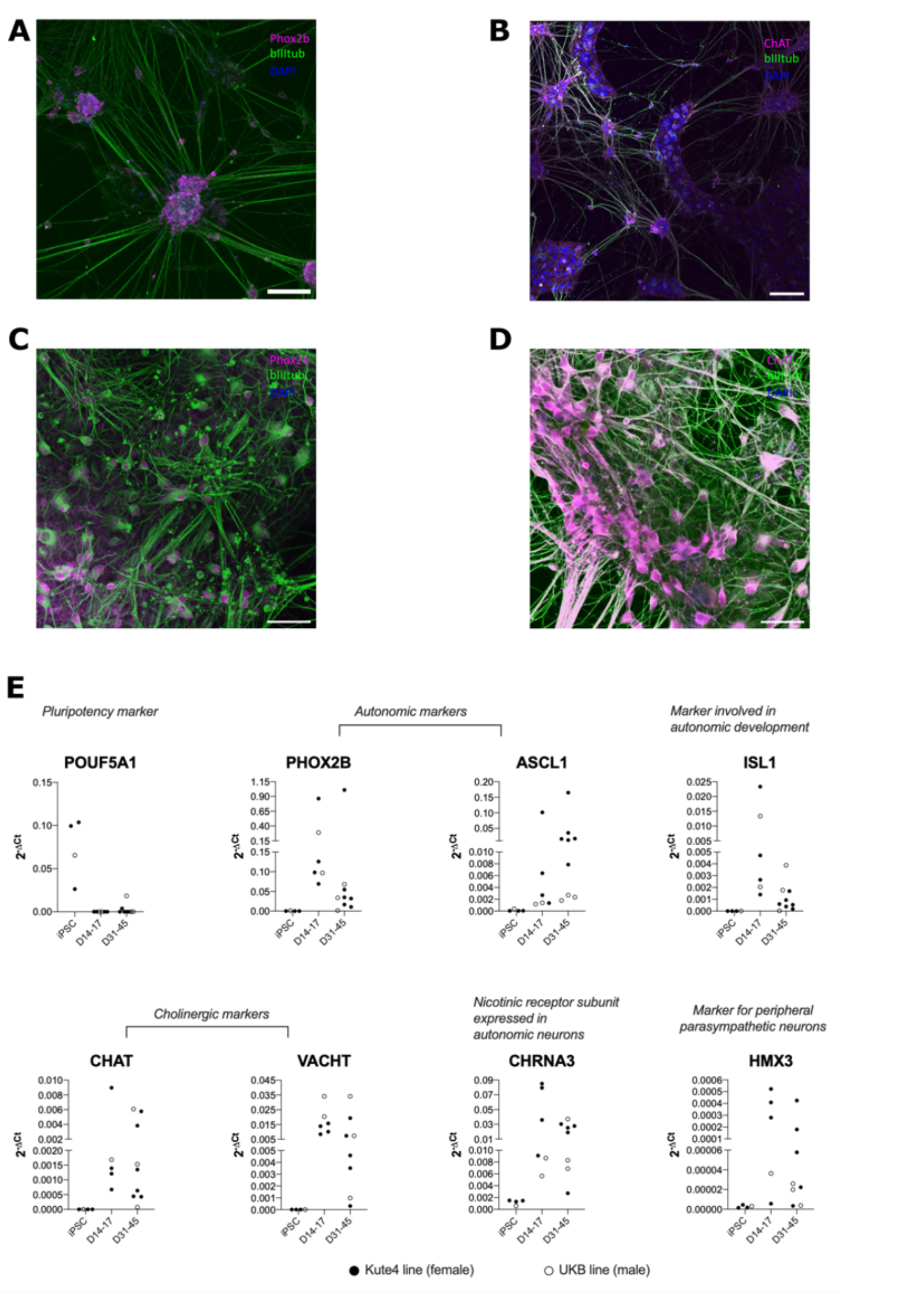
Investigation of neuronal identity using a combination of immunocytochemistry (A-D) and RT-qPCR (E). A and C Phox2b, beta 3 tubulin, DAPI. B and D, CHAT, beta 3 tubulin and DAPI. Scale bar in A and B 100*μ*m; scale bar in C and D 50*μ*m.Of note, the neurons start expressing autonomic and cholinergic markers from Day 14. (E) RT-qPCR of the following markers: *POUF5A1, PHOX2B, ASCL1, ISL1, CHAT, VACHT, CHRNA3* and *HMX3*. Each dot represents one distinct differentiation batch, filled circles Kute4 line, empty circles UKB line.

**NOTE:** By day 14-17 cells should already express the autonomic markers *ASCL1* and *PHOX2B* together with the cholinergic markers *CHAT* and *VACHT*.

**CRITICAL:** Combining 2-3 coverslips per sample ensures enough RNA for RT-qPCR with the RNeasy Plus Micro kit. Other methods (e.g. Smart-seq2^5^) could allow the use of less starting material (e.g. one coverslip) for RNA isolation.

### Functional assessment: whole-cell patch clamp

#### Timing: 4hours

This section describes the employment of whole-cell patch clamp recordings to analyze the electrophysiological properties of the neurons and their action potential profile. As shown in Figure 3, the resting membrane potential becomes more hyperpolarized (i.e. more negative) over time in culture, consistent with a more mature-like profile from day 39 compared to earlier stages of differentiation. Using this technique we also found that our hiPSC-derived parasympathetic neurons can be sub-grouped into phasic and tonic consistent with native parasympathetic neurons of e.g. intracardiac ganglia^6^.

**Figure 3.**
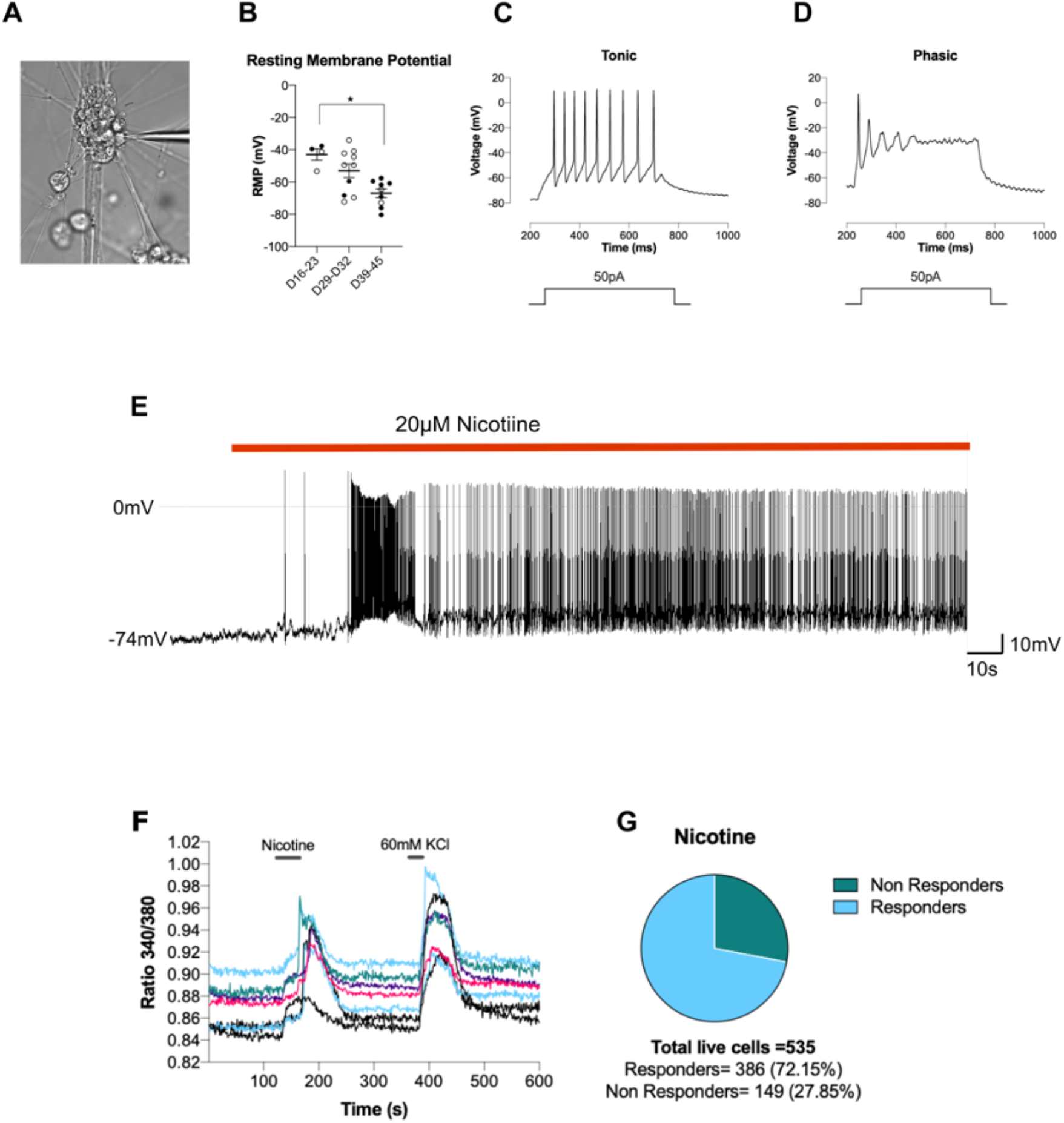
Functional characterization of hiPSC-parasympathetic neurons using whole cell patch clamp recordings (A-E) and Ca^2+^ imaging with Fura-2 (F). (A) Phase contrast image of a patched neuron. (B) Analyses of resting membrane potential from Day 16. The neurons have a more hyperpolarized resting membrane potential from >Day 39, suggesting a more mature phenotype. Each dot represent a single cell. Filled circle Kute4, empty circles UKB line. Kruskal-Wallis test with p=0.0087; Dunn’s multiple comparisons test p=0.0106 between D19-23 and D39-45. At least two differentiation per line per timepoint. In total three batches for the Kute4 line and 2 for the UKB line were used. (C) Sample trace of a tonic (D) and a phasic neuron. (E) Nicotine (20*μ*M) application elicits action potentials in a hiPSC-parasympathetic neuron (whole cell patch clamp current clamp recording at resting membrane potential in gap free mode). (F-G) Ca^2+^ imaging using Fura-2. (F) Sample traces showing responses following 10*μ*M nicotine and 60mM KCl. (G) Percentage of nicotine responders among the live excitable cells from one coverslip of Day 39 Kute4 hiPSC-parasympathetic neurons.

15. Whole-cell patch clamp electrophysiology
  a. Pull pipettes from glass capillaries using a 2-step puller, aiming for a 4.5-6MΩ resistance
  b. Gently transfer the coverslip with the neurons to the bath recording chamber in a patch clamp electrophysiology setup with the external solution (see Table 6 for recipe of the one we used)
  c. Fill the pipette with intracellular solution (Table 7 illustrates the one we used)
  d. Gently place the pipette in the bath solution, apply some positive pressure, approach the cell of interest and form a Giga Ohm seal using negative pressure
  e. Apply more negative pressure to achieve the whole-cell configuration
  f. Switch to current clamp mode to measure resting membrane potential and investigate the firing patterns of the neurons. Evoked action potentials were recorded at a 0.5 Hz frequency using a 500-millisecond current injection

**Table 6.** External solution employed for whole-cell patch clamp electrophysiology (bubbled with 95% O_2_ and 5% CO_2_)

|  | (mM) |
| --- | --- |
| NaCl | 124 |
| KCl | 2.5 |
| NaHCO<sub>3</sub> | 26 |
| NaH<sub>2</sub>PO<sub>4</sub> | 1 |
| CaCl<sub>2</sub> | 2 |
| MgCl<sub>2</sub> | 1 |
| Glucose | 10 |
| pH 7.4 |  |

**Table 7.**
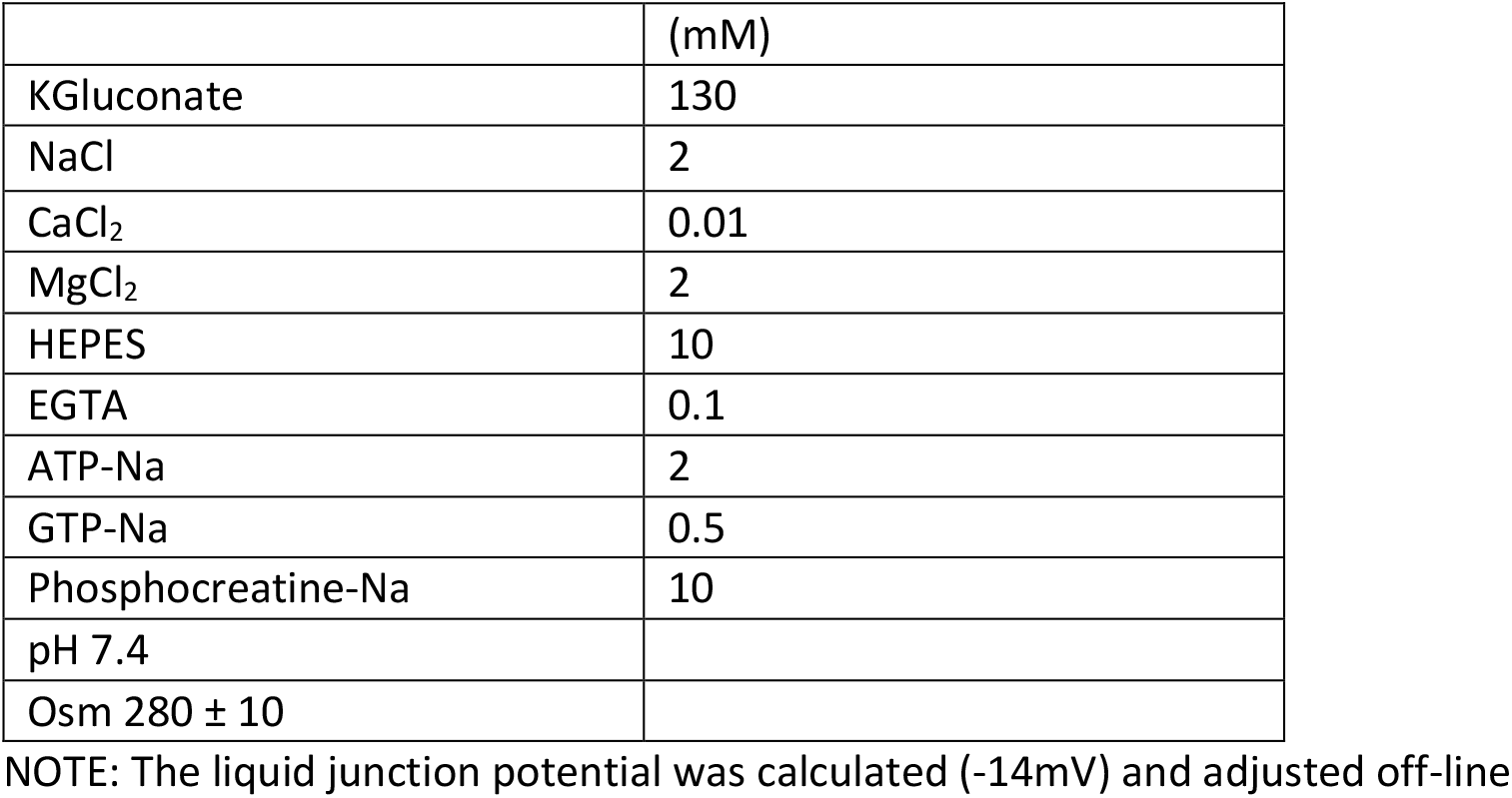
Internal solution employed for whole-cell patch clamp electrophysiology.

**CRITICAL:** If performing whole-cell patch clamp recordings, Geltrex supplementation can form a sticky layer on top of the cells and make it difficult to access cells with a patch pipette. In this case, perform media changes without Geltrex on the designated media change days just before the recording.

**NOTE:** There are numerous critical and troubleshooting steps for whole-cell patch clamp recordings, which are beyond the scope of this manuscript. Some useful resources include^7,8^, which can be applied for multiple cell types.

### Functional assessment-Ca^2+^ imaging

#### Timing: 2hours

This section shows how to assess the response of hiPSC-derived parasympathetic neurons to nicotine using Ca^2+^ imaging with ratiometric Ca^2+^ indicators; we used the Fura-2 dye.

16. Ca^2+^ imaging (Fura-2 dye)
  a. Incubate a coverslip (37°C, 5%CO_2_) with 500*μ*L of Fura-2 (2.5*μ*g/mL Fura2AM, 0.01% Pluronic acid, 10mM probenecid) in Extracellular Solution (Table 8)
  b. Place the coverslip in the recording chamber with perfusion of extracellular solution
  c. Perform Ca^2+^ imaging. We employed a 10x objective and obtained the measurements at 340nm and at 380nm. We performed a brief application of nicotine (10*μ*M) to test if the cells had functional nicotinic receptors and 60mM KCL (Table 9) to identify live (and excitable) cells
  d. The image stack of the recordings for both 340nm and 380nm were saved as tiff
17. Calcium imaging analyses
  a. Open the 340nm and 380nm files, convert them into 32-bit
  b. Run the Fiji Plugin Ratio Plus, Image A (340nm Numerator), Image B (380nm Denominator). The Region of interests (ROI) were identified using the Fiji Plugin StarDist followed by manual inspection. A positive response to 60mM KCl or nicotine was identified as: more than 10 frames had to exceed the respective baseline by at least 3 standard deviations, as described in our previous work^9^

**Table 8:** External solution for Fura-2 Ca^2+^ imaging (neurons alone)

|  | In mM |
| --- | --- |
| NaCl | 140 |
| KCl | 5 |
| CaCl<sub>2</sub> | 2 |
| MgCl<sub>2</sub> | 1 |
| HEPES | 10 |
| Glucose | 10 |
| PH 7.4 |  |

**Table 9:** 60mM solution for Fura-2 Ca^2+^ imaging (neurons alone)

|  | In mM |
| --- | --- |
| NaCl | 85 |
| KCl | 60 |
| CaCl<sub>2</sub> | 2 |
| MgCl<sub>2</sub> | 1 |
| HEPES | 10 |
| Glucose | 10 |
| PH 7.4 |  |

### Differentiation of hiPSC cells into atrial cardiomyocytes

This section explains how we differentiated atrial-like cardiomyocytes from hiPSC. We adapted the protocol from^10^ using the HPSI0614i-ciwj_2 line. We were interested in employing atrial-like cardiomyocytes to mimic the high innervation of the atria by parasympathetic neurons.

#### Timing: 20-30 days

18. Differentiation of hiPSC-derived atrial cardiomyocytes
  a. Plate hiPSC (cultured and passaged as described in numbers 3-5) onto vitronectin coated wells (30 wells of a 48 well plate from 2 wells of a 6 well plate)
  b. Differentiation is initiated by adding Day 0 media (Table 10) when cells are 90% confluent
  c. Media is changed as detailed in Table 6 on Days 2, 3, 4 and 6
  d. From day 8 onwards, media can be changed every 2-3 days

**Table 10:** hiPSC-atrial cardiomyocytes media. The base media for each day is RPMI 1640 + Glutamax, 25mM HEPES, 1% Pen/Strep with additional supplements as detailed.

| Day | Concentration | Amount (15mL) |
| --- | --- | --- |
| 0 | 4μM CHIR99021<br>7.5μM Albumin<br>750μM Ascorbic Acid | 6μL CHIR99021<br>100μL Albumin<br>50μL Ascorbic Acid |
| 2 | 5μM IWP2<br>7.5μM Albumin<br>750μM Ascorbic Acid | 15μL IWP2<br>100μL Albumin<br>50μL Ascorbic Acid |
| 3 | 5μM IWP2<br>1μM Retinoic Acid<br>7.5μM Albumin<br>750μM Ascorbic Acid | 15μL IWP2<br>5μL Retinoic Acid<br>100μL Albumin<br>50μL Ascorbic Acid |
| 4 | 1 μM Retinoic Acid<br>7.5μM Albumin<br>750μM Ascorbic Acid | 5μL Retinoic Acid<br>100μL Albumin<br>50μL Ascorbic Acid |
| 6 | 7.5μM Albumin<br>750μM Ascorbic Acid | 100μL Albumin<br>50μL Ascorbic Acid |
| 8+ (change media every 2-3 days) | 1X B27 | 300μL B27 |

**CRITICAL:** Spontaneously beating regions are visible as early as Days 7-10. Cells which didn’t show spontaneous beating after Day 12 were discarded.

19. Quality control for hiPSC-derived atrial cardiomyocytes:
  a. Collect cells for RNA isolation using the RNeasy Plus Micro kit, 1 to 2 wells of 24 or 48 well plates generate enough RNA for analysis
  b. Convert RNA to cDNA using High-capacity cDNA reverse transcriptase kit Perform RT-qPCR to check gene expression of key markers such as TNNT2, MYL2, MYL7 using Taqman probes (Figure 4)
  c. Fix coverslips > Day 21 with 50% Methanol/50% Acetic acid on ice for 10 mins for antibody staining with TNNT2, MYL2, MYL7

**Figure 4.**
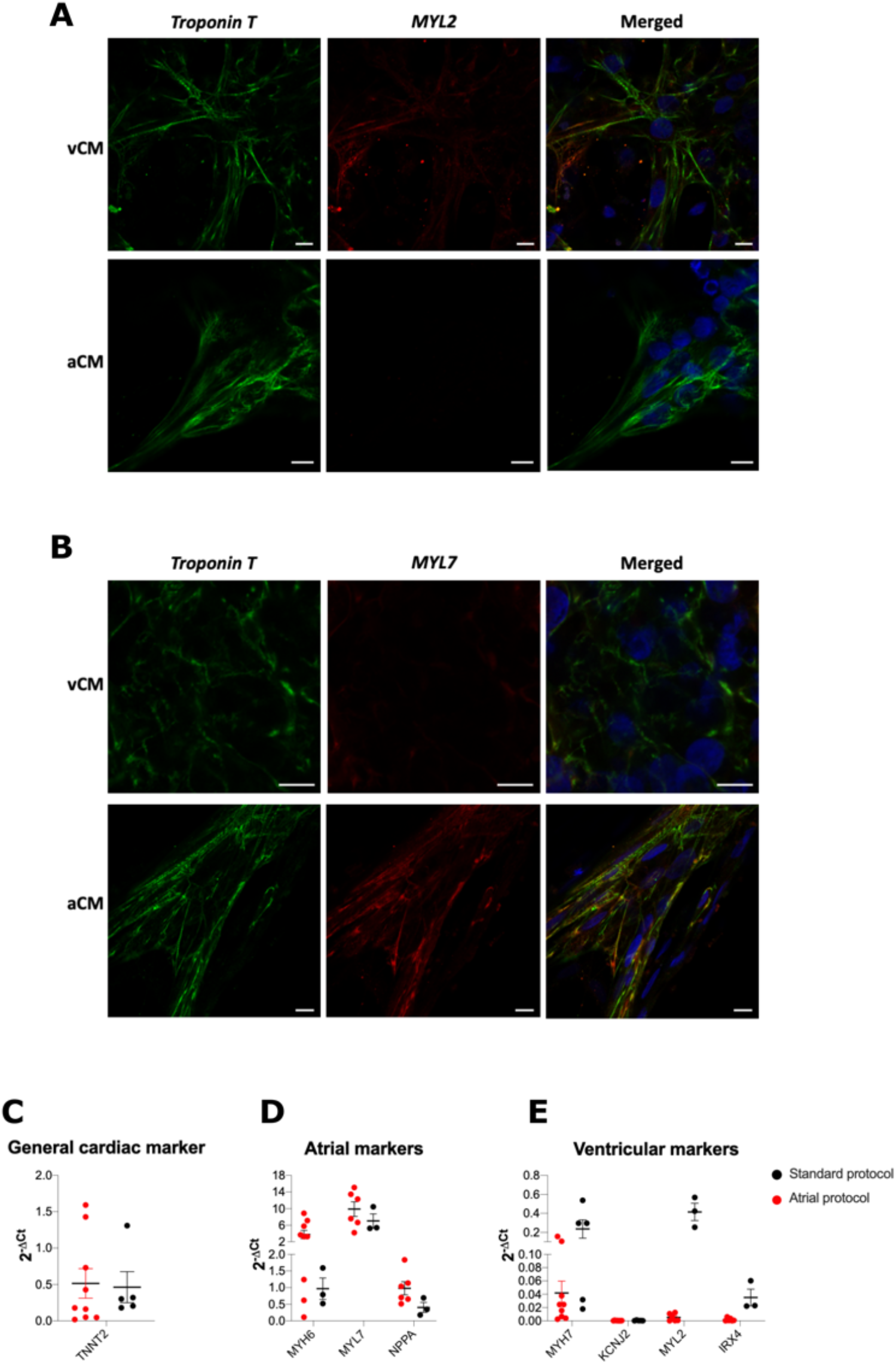
Characterization of hiPSC-derived cardiomyocytes using the atrial protocol^10^ (aCM) and the standard ventricular protocol^10^ (vCM). (A) Immunocytochemistry for the ventricular marker MYL2. (B) Immunocytochemistry for the atrial marker MYL7. Taqman analyses for general cardiac markers (C), atrial markers (D) and ventricular markers (E). Each dot represents a distinct cardiomyocyte differentiation.

**NOTE:** We further confirmed the atrial lineage by differentiating, in parallel, hiPSC into cardiomyocytes with a ventricular like phenotype using the same protocol but removing retinoic acid as in^10^ incubation between Days 3-6 (Figure 4). Atrial markers, *MYL7*, in hiPSC-derived atrial cardiomyocytes were present by Day 21 when RNA was harvested to assess levels of gene expression via qPCR. Cardiomyocyte marker, *TNNT2*, was also assessed as well as ventricular marker, *MYL2*.

### Coculture of hiPSC-derived parasympathetic neurons and hiPSC-derived atrial cardiomyocytes

This section explains how we set up a 2D coculture system that could be employed to study the interaction between parasympathetic neurons and atrial cardiomyocyte in physiological conditions and disease states. Dysfunction of parasympathetic neurons contributes to diseases like atrial fibrillation.

#### Timing: 7 days

20. Coculture of hiPSC-derived atrial cardiomyocytes and hiPSC-derived parasympathetic neurons
  a. Dissociate cardiomyocytes with Tryple (10min @37°C)
  b. Centrifuge @ 280g for 5 min
  c. Remove supernatant
  d. Resuspend, using gentle pipetting, three wells from a 48 well plate of iPSC-cardiomyocytes (or 1 well of a 24 well plate) into 1.5mL co-culture media (Table 11) and distribute in 3 coverslips/wells (with or without neurons). 500*μ*L of cardiomyocytes were employed per coverslip
  e. Remove neuronal media from mature neurons (we used Day 42) and plate cardiomyocytes very carefully as if you were changing media without disrupting the neurons, i.e. plating on the side and sometimes gently rotating the well to allow even distribution
  f. Plate cardiomyocytes in monoculture employing the same co-culture media on Geltrex coated coverslips
  g. Maintain the co-culture for a week with co-culture media, changing media every 2-3 days

**Table 11:**
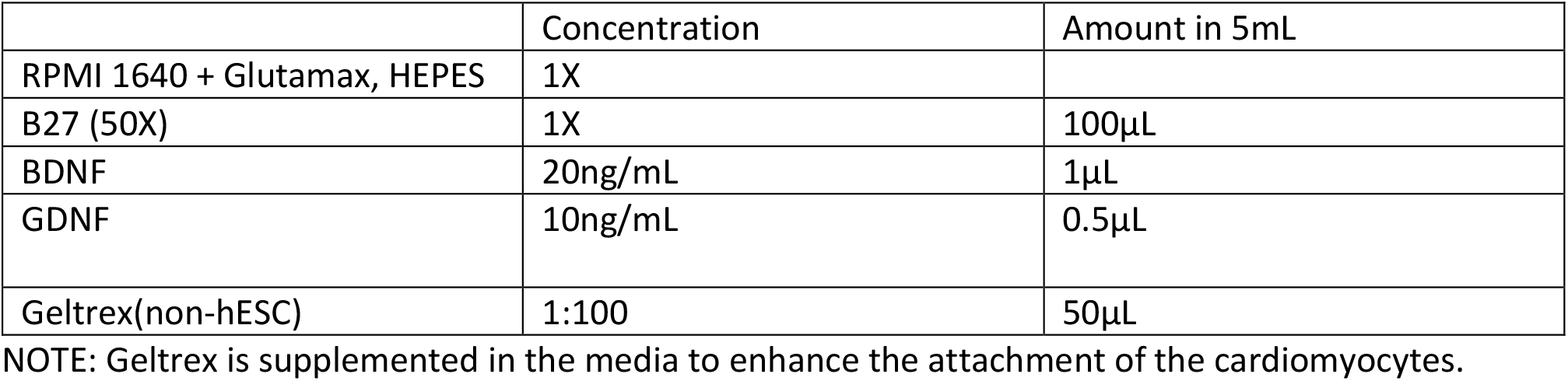
Coculture media.

**Table 12:** Co-culture recording solution.

|  | (mM) |
| --- | --- |
| NaCl | 130 |
| KCl | 5 |
| MgCl | 0.4 |
| CaCl | 0.4 |
| Glucose | 10 |
| HEPES | 10 |
| pH 7.4 (with NaOH) |  |

**CRITICAL:** Be gentle when resuspending hiPSC-cardiomyocytes, pipette up and down 5 times.

### Functional assessment of the coculture-Ca^2+^ imaging

We assessed the functional interaction between hiPSC-derived parasympathetic neurons and hiPSC-atrial cardiomyocytes following seven days of coculture. The success of co-culturing the two cell types is evident by the immunocytochemistry (Figure 5A) and the reduction of spontaneous Ca^2+^ transients of the cardiomyocytes following nicotine (20*μ*M) application (Figure 5B-E).

**Figure 5.**
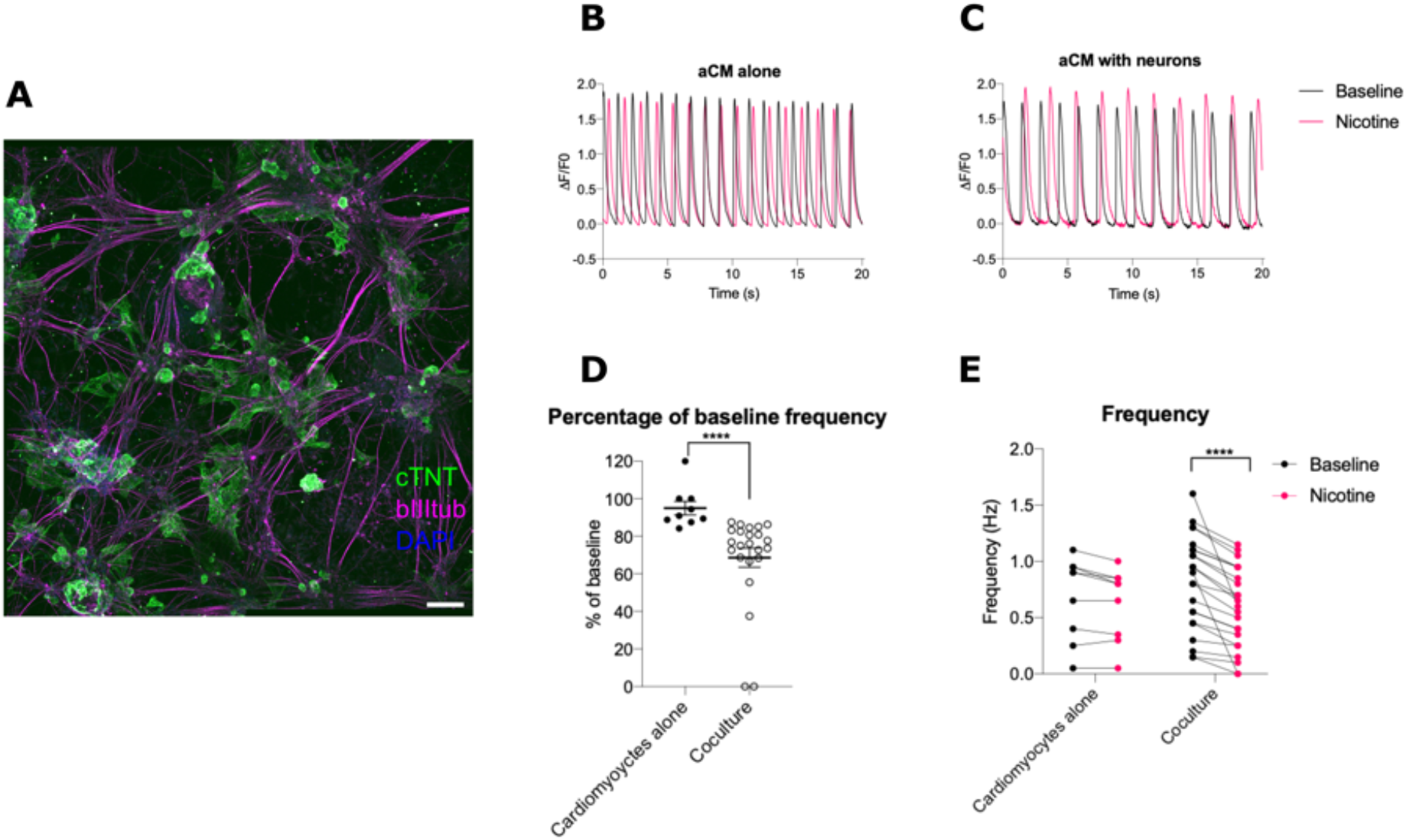
Coculture of hiPSC-parasympathetic neurons and hiPSC-atrial cardiomyocytes. (A) Immunocytochemistry for a cardiac marker (cardiac troponin) and a neuronal marker (beta III tubulin). (B) Sample traces of spontaneous Ca^2+^ transients (using Fluo-4) of iPSC-cardiomyocytes alone and cardiomyocytes co-cultured with iPSC-parasympathetic neurons (C) at baseline (black) and in the presence of nicotine (magenta). (D) Change in spontaneous Ca^2+^ transients as a percentage from baseline following nicotine application. Mann Whitney test p<0.0001. (E) Frequency of spontaneous Ca^2+^ transients before and after nicotine application. Two-way ANOVA p<0.0001, with Sidak’s multiple comparison test: cardiomyocyte baseline vs nicotine p>0.05; coculture baseline vs nicotine p<0.0001. Each dot represents distinct regions of rhythmic spontaneous Ca^2+^ fluctuations identified as ROIs in Fiji; one cannot be sure they are single cardiomyocytes. 1 differentiation of cardiomyocytes, 1 differentiation of neurons (D51); n=9 cardiomyocyte ROIs alone (2 coverslips), n=23 coculture ROIs (3 coverslips).

**Figure 6.**
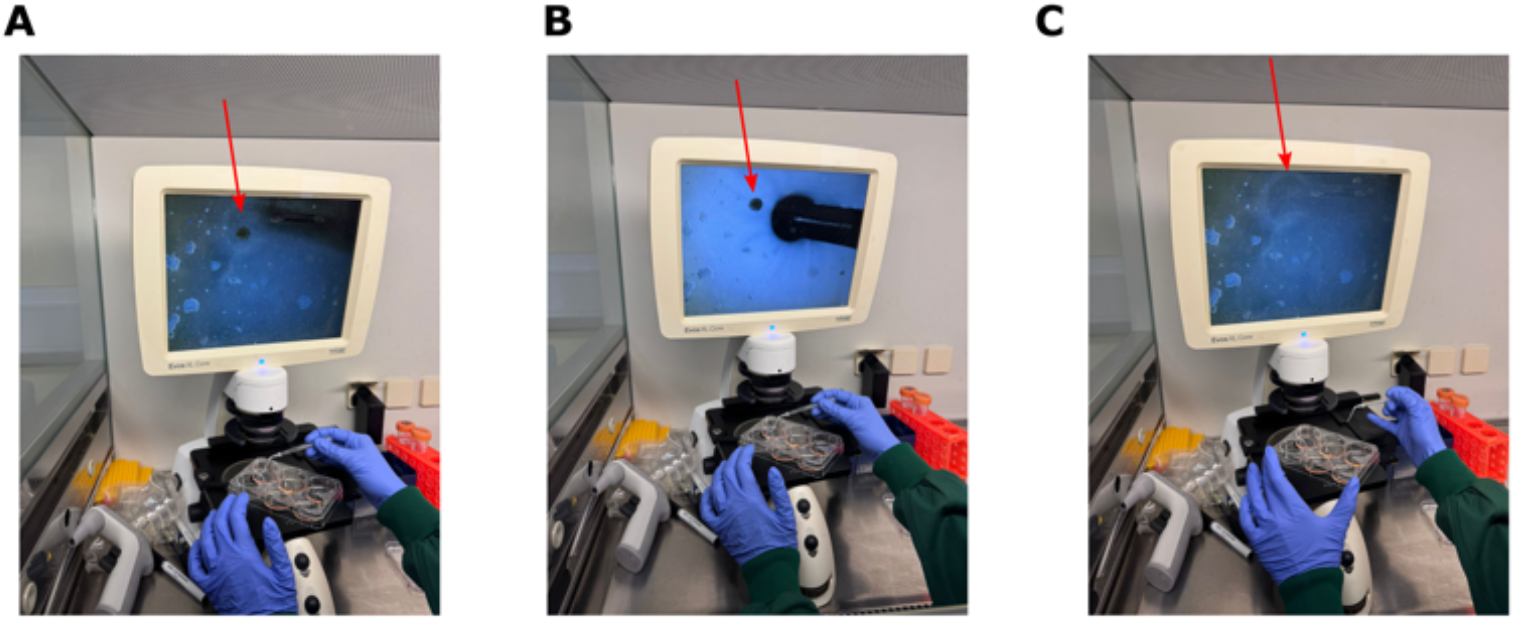
Removal of hiPSC grown in 3D. Arrow point at a bad looking hiPSC colony grown in 3D (A and B) and removed in C. A P10 tip was bent and attached to a P1000 tip. As shown in B it was carefully approached to the 3D grown hiPSC colony. C shows the complete removal of the colony.

#### Timing: 1 hour

21. Ca^2+^ imaging
  a. Incubate the cells with 1*μ*M Fluo4-AM in coculture recording solution (Table 12) for 20 minutes
  b. Wash cells with recording solution once
  c. Cells are ready for Ca^2+^ imaging. We recorded spontaneous Ca^2+^ transients for 20 seconds at baseline and following nicotine application (after 1 min of drug application).
22. Ca^2+^ imaging analyses:
  a. Import videos as tiff files
  b. Identify the region of spontaneous Ca^2+^ fluctuation using the standard deviation function (Image Stack>Z project> standard deviation) in the baseline videos. The spontaneous Ca^2+^ transient frequency was assessed by counting the number of peaks. The same regions of interests were employed following nicotine application

### Expected outcomes

Here, we describe a protocol to differentiate hiPSCs into parasympathetic neurons that have functional and electrophysiological properties similar to the native ones. To ensure the reproducibility of the protocol we employed two distinct hiPSC lines, one male and one female, which also required distinct extracellular matrices for maintenance. The first checkpoint for efficient differentiation is the formation of smooth spheroids in the embryoid body stage (day 0-11) and neuronal-like projection as early as day 14 (Figure 1 and see Troubleshooting problem 3). The second checkpoint is the verification of their expression (RT-qPCR) at Day 14-17 of autonomic (*ASCL1, PHOX2B*) and parasympathetic (*CHAT, VACHT*) markers, as well as markers involved in autonomic development (e.g. ISL1, Figure 2). At a more mature stage (>Day 30) they should preserve the expression of autonomic and parasympathetic markers using RT-qPCR and immunocytochemistry (Figure 2). We also assessed for their expression of the postganglionic parasympathetic marker *HMX3*^3^, not expressed in undifferentiated hiPSC, suggesting some postganglionic lineage.

A recent systematic review^3^ on stem cell derived autonomic neurons highlighted the importance of functional assessments of differentiated cells for quality control, as well as co-culture with other cell types. Our functional assessment (Figure 3B) of hiPSC-parasympathetic neurons revealed that from D39, hiPSC-parasympathetic neurons have a resting membrane potential (averaged -65mV) comparable to native counterparts, suggesting a mature-like phenotype. It also showed that the neurons can be classified into phasic and tonic firing (Figure 3C-D), and that they responded to nicotine, shown both using whole-cell patch clamp recordings (Figure 3E) and Ca^2+^ imaging (Figure 3F). As highlighted by Bos et al^3^ and James et al^2^ the parasympathetic system is critical in controlling target tissue, we therefore developed a coculture system with hiPSC-derived atrial cardiomyocytes to mimic their key role in controlling atrial electrophysiology^11^. We first differentiated hiPSCs into atrial cardiomyocytes using retinoic acid at Day 3-6 as per^10^. We verified their atrial-like profile (Figure 4) comparing them with hiPSC differentiated into cardiomyocytes without retinoic acid, therefore more ventricular-like. Establishing the coculture (Figure 5) of the two cell types, revealed the functional modulation of hiPSC-atrial cardiomyocytes by hiPSC-parasympathetic neurons: nicotine reduced their spontaneous activity (Figure 5B-E, Video 1).

## Limitations

Using hiPSC-derived parasympathetic neurons in research offers significant advantages, among them, they are human and highly scalable. Parasympathetic ganglia lie adjacent to their target organs, making them very difficult to access for primary culture, with low number of neurons per animal. hiPSC-parasympathetic neurons are therefore ideal to use for drug discovery studies, and our neurocardiac model would align with the recent move of pharmaceutical companies (e.g. GSK) to employ hiPSC-based cardiac models for primary cardiotoxicity screens^12^. Generating hiPSCs from patient cells and their isogenic controls can also allow the investigation of the impact of genetic profile on cellular physiology.

A few technical steps should be considered to ensure the success of the differentiation. The main limiting factors of our hiPSC-derived parasympathetic neuron protocol are: healthy hiPSC colonies (see Troubleshooting, Problem 1) and using a good quality of hiPSC line (e.g. healthy and pluripotent). As shown in Figure 8 and described in Problem 3, a substandard (low pluripotent) hiPSC line resulted in unsuccessful differentiation.

Moreover, as shown in Figure 1B, when initiating the protocol (Day 0) hiPSC colonies should look healthy (see Troubleshooting 1). Starting with distinct hiPSC colonies will allow the smooth formation of embryoid bodies when detached with collagenase. Finally, appropriate coating of the coverslip with Geltrex is critical for the success of the protocol: poor coating can result in cell death and lack of neurite outgrowth (See Problem 2).

Because parasympathetic neurons lie very close to their target organs, they closely interact with their surrounding cells. Using parasympathetic neurons in isolation might therefore not fully reproduce their physiological function. We overcame this with our coculture system between hiPSC-parasympathetic neurons and hiPSC-atrial cardiomyocytes (Figure 5). A similar approach was used by Wu et al^4^ who co-cultured hiPSC-parasympathetic neurons and hiPSC-cardiomyocytes (protocol for mixed types of cardiomyocytes). Another approach to consider could be to incorporate hiPSC-parasympathetic neurons into sympathetic innervated heart assembloids recently developed by Zeltner et al^13^ or to incorporate other cells that might be important for the research question, e.g. immune cells.

It is also important to note that hiPSC-derived cells are unlikely to reproduce the full transcriptomic range and diversity of their native counterparts. Here, we tested key genes and key functional properties of parasympathetic neurons, showing that our protocol generates physiological features of parasympathetic neurons. However, we would advise testing for the expression of genes relevant to a particular research question before starting any experiments. We tested for the *HMX3* gene, suggesting some postganglionic lineage; preganglionic-like neurons might also be present. There might also be other cells in the dish, with their numbers likely to vary with differentiation efficiency; more prolonged time in culture might require treatment with Cytosine Arabinoside or similar drugs to remove proliferating non-neuronal cells

## Troubleshooting

### Problem 1

Some hiPSCs do not look healthy

### Potential solution

If some cells grow in 3D or there are some small areas of spontaneous differentiation, remove the 3D colonies/spontaneously differentiated cells with a bent p10 tip (Figure 6). This will ensure the employment of only healthy colonies.

### Problem 2

Cells die following plating or do not form neurites.

### Potential solution

Poor cell plating/cell death or lack of neurite formation might occur if cells are not seeded on Geltrex coated areas or if Geltrex is not fully removed or dries out before cell seeding. We coat coverslips with Geltrex just before or during the cell dissociation step to make sure there are less than 2hours (ideally 1h) between coating and cell seeding. We illustrate optimal vs inadequate coverslip coating with Geltrex in Figure 7. The top two wells in Figure 7a illustrate the optimal placement of the Geltrex bubble, the bottom the spreading of the bubble which would not allow adequate attachment of cells. Figure 7b shows the removal of Geltrex coating just before seeding. The top wells show the optimal residue of Geltrex just before cell seeding, the bottom two poor coating which will result in poor cell survival/neurite growth.

**Figure 7.**
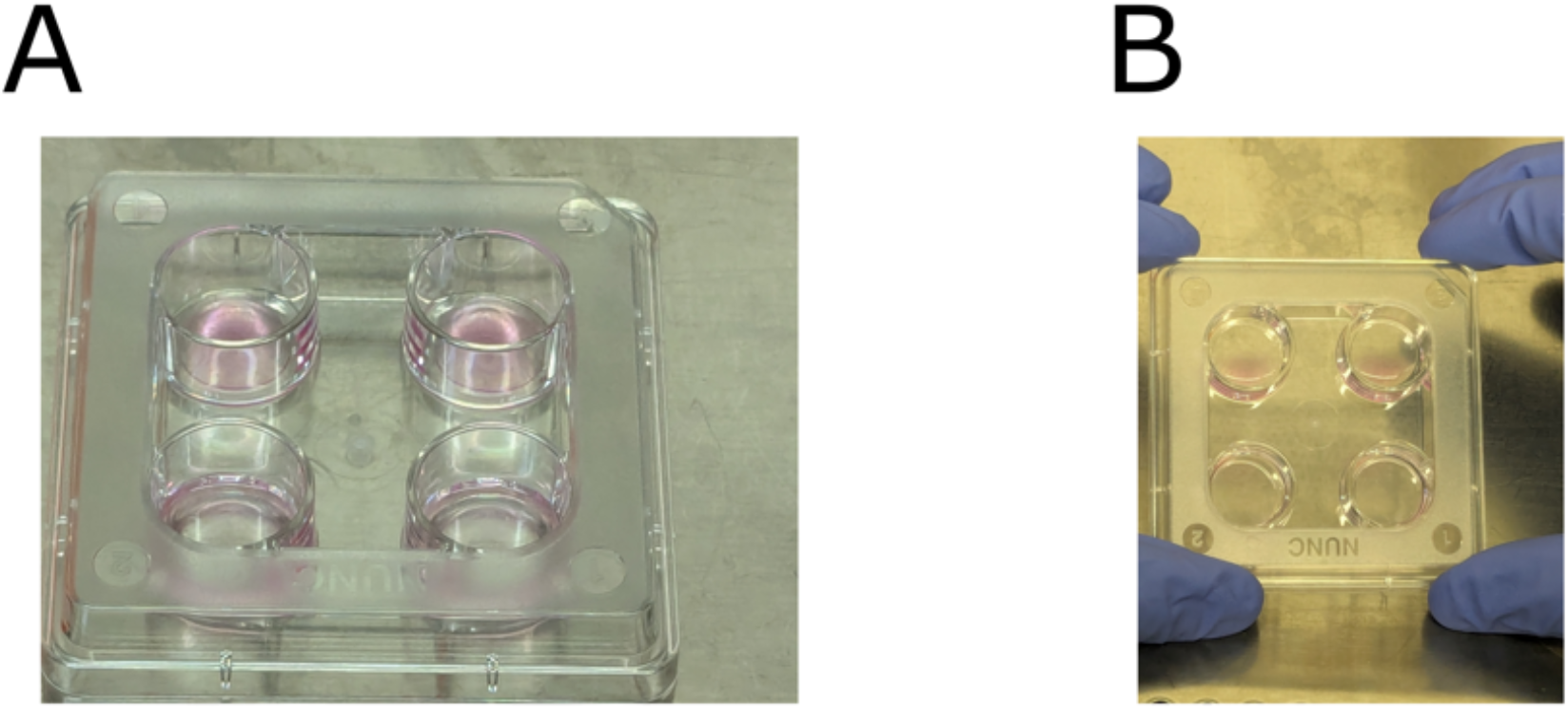
Appropriate versus inadequate coating of coverslips with Geltrex. (A) Top wells: optimal coating of the coverslip with Geltrex solution, as evidenced by the formation of bubbles. Bottom wells: Geltrex spread across the coverslips without forming a bubble. (B) Top wells: optimal residue of Geltrex just before cells seeding. Bottom wells: the removal of the Geltrex solution following the lack of formation of the Geltrex bubble resulted in inappropriate coverslip coating which will likely result in poor survival/neurite growth.

**Figure 8.**
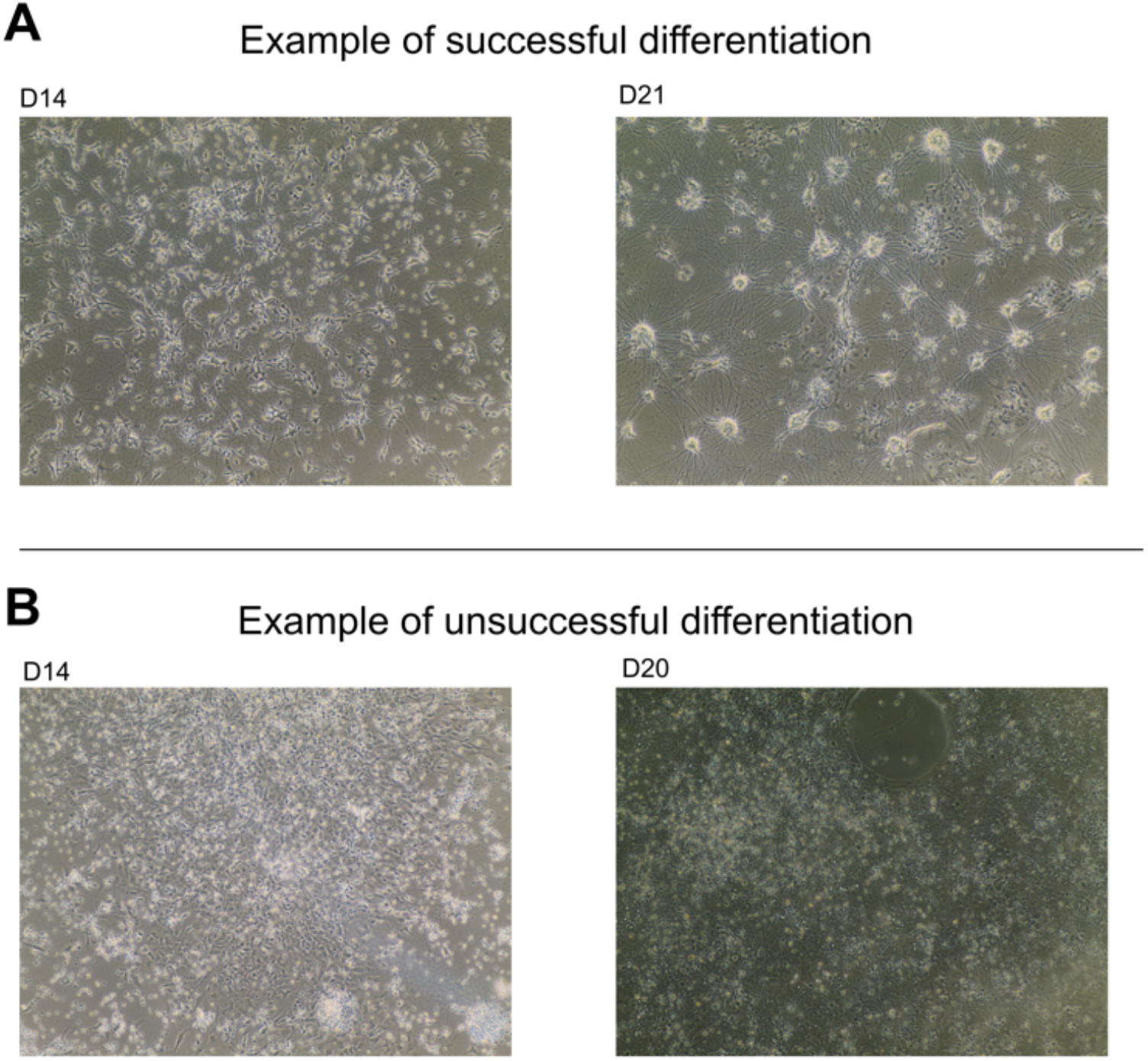
Unsuccessful differentiation due to substandard hiPSC line. (A) Example of successful generation of neurons from a ‘good’ hiPSC line at Day 14 and Day 21, respectively. (B) Unsuccessful generation of neurites from a ‘bad’ hiPSC line at Day 14 and Day 20.

### Problem 3

Unsuccessful generation of neuronal like cells at Day 14.

### Potential solution

Verify that the original hiPSC line meets the standard. A substandard, e.g. low pluripotent line, can result in unsuccessful generation of neurons; we experienced exactly this problem, which we traced back to a problem with hiPSC line quality. Figure 8A shows a successful generation of neurons at Day 14 and 21 compared to the unsuccessful generation of neurons (Figure 8B) evident by the lack of neurites at both Day 14 and 21.

## Resource availability

## Materials and equipment

We have employed commercially available reagents. The hiPSC lines Kute4 and HPSI0614i-ciwj_2 can be obtained from the HipSci website.

## Acknowledgments

This work was supported by an Early Career Research Award IoPPN (Institute of Psychiatry, Psychology and Neuroscience) King’s College London to LF. F. Denk has received support from the UKRI Strategic Priorities Fund (SPF) Advanced Pain Discovery Platform (APDP, MR/W027518/1). J.F.N. acknowledges support from The Royal Society (RGS\R2\202265) and The Lister Institute of Preventive Medicine. A.M.T. and A.T acknowledge support from the NIHR Barts Biomedical Research Centre. I.N.C was funded by the Wellcome Trust as part of the “Neuro-Immune Interactions in Health & Disease” Wellcome Trust PhD Program (218452/Z/19/Z).

We acknowledge Life & Brain GmbH in Bonn and the University Hospital of Bonn for provision of the UKBi013 iPSC line. We thank Professor J.F.Brunet for the Phox2b antibody.

## Author contributions

L.F. provided funding, designed the experiments, differentiated the cells into hiPSC-parasympathetic neurons, performed all the hiPSC-parasympathetic neurons experiments, optimized the protocol, analyzed the data and wrote the manuscript; A.M.T. differentiated the cells into hiPSC-cardiomyocytes and characterized the cardiomyocytes. L.F and A.M.T. designed and established the co-culture and performed the experiments; I.N.C. and J.F.N. helped in the design and optimization of the hiPSC-parasympathetic protocol; A.T. provided the funding and helped designing the cardiomyocytes and coculture experiments; F.D. provided funding and helped designing the experiments. All authors reviewed the manuscript

## Figure legends

Video 1. Example video showing spontaneous activity (Ca^2+^) of the cardiomyocytes when in coculture with hiPSC-parasympathetic neurons.

## Key resources table

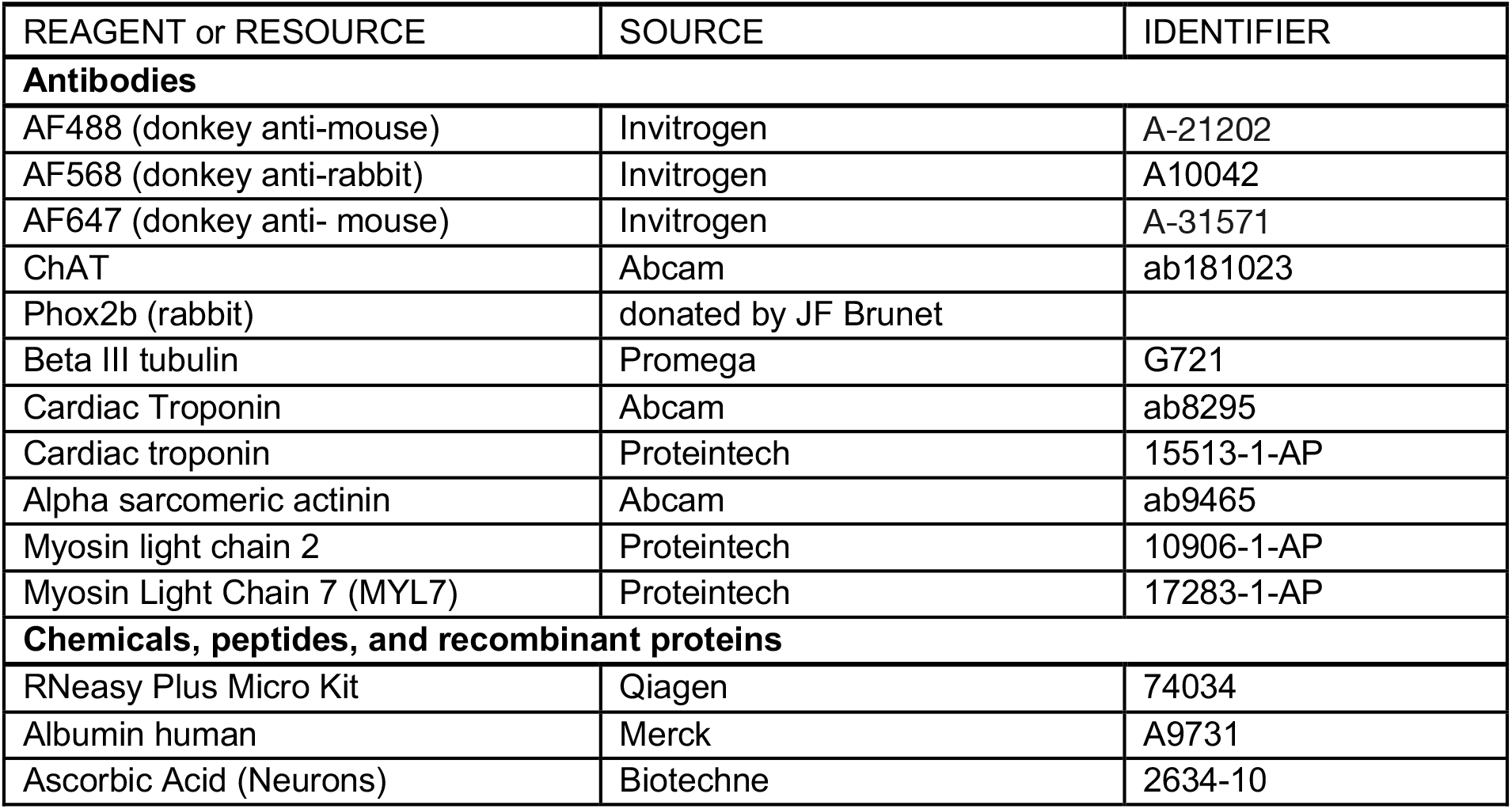
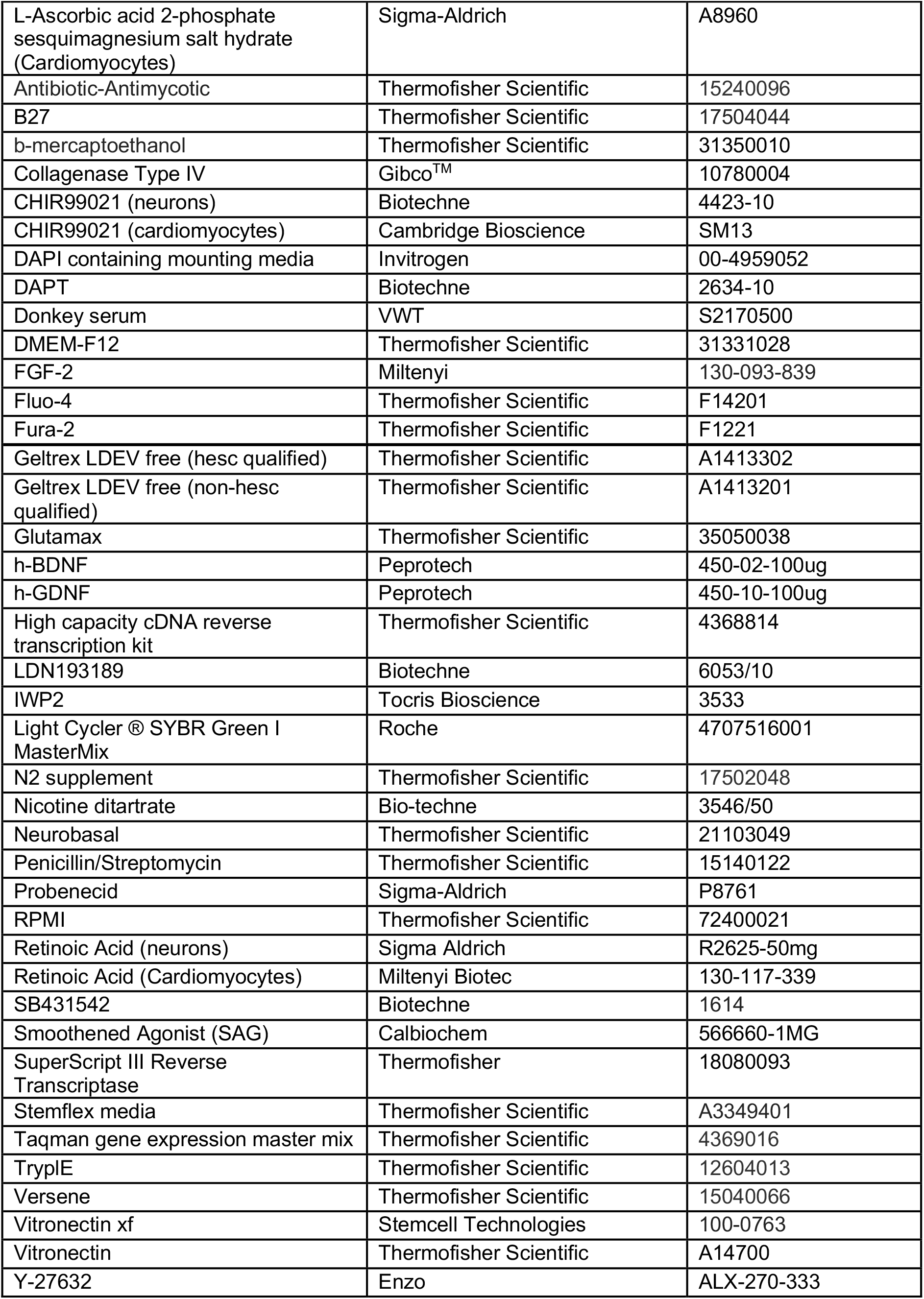
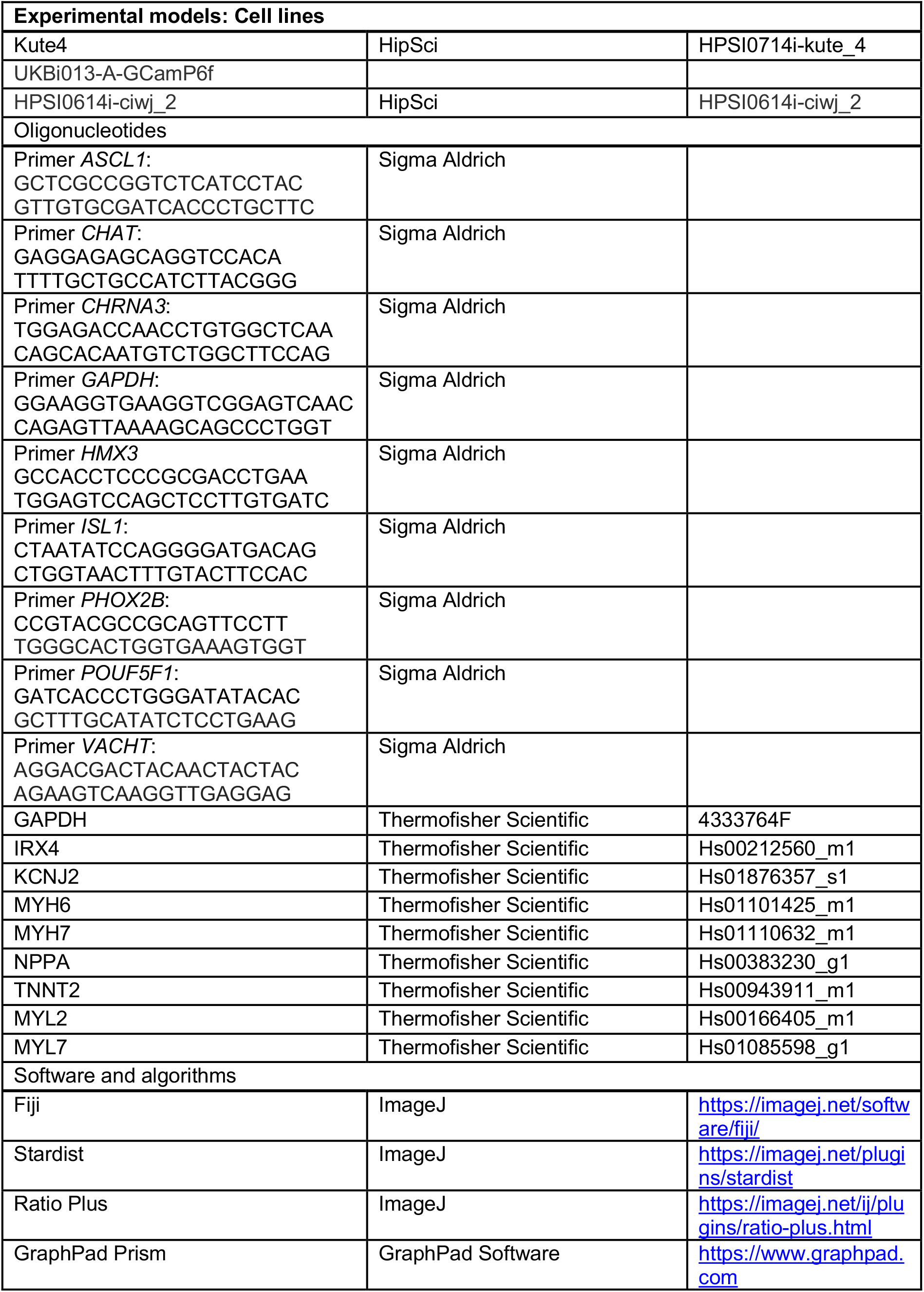
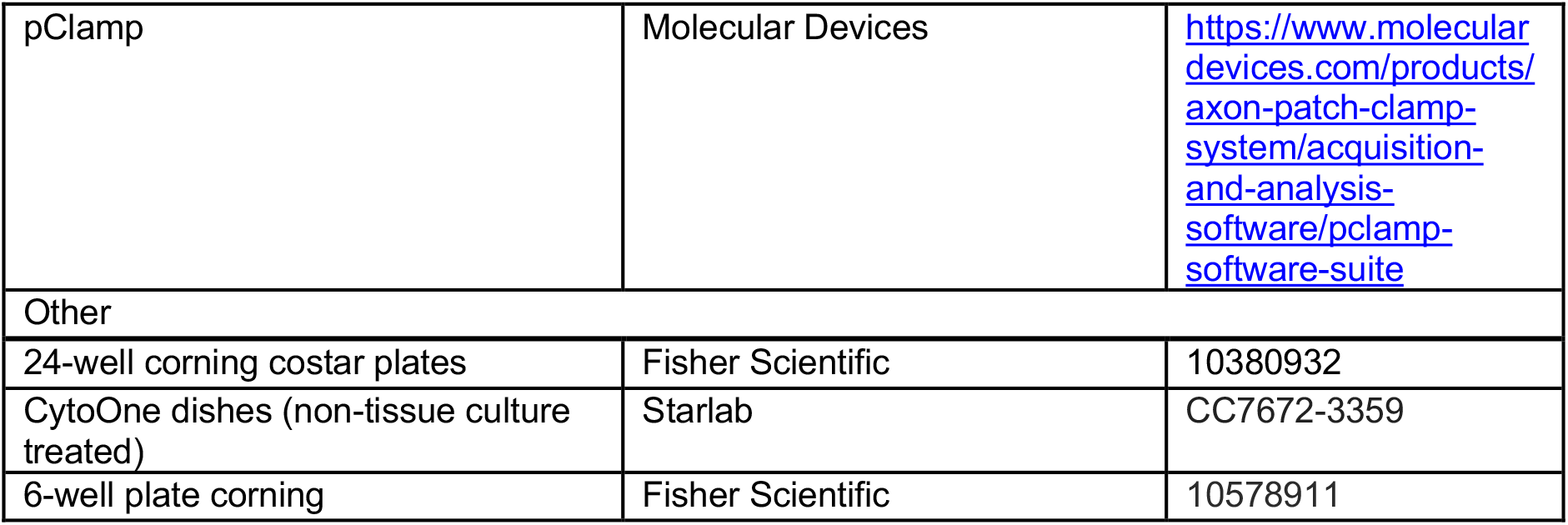

